# TP53 Intron Derived Concentrations Implicate p53

**DOI:** 10.1101/307959

**Authors:** Kevin Bermeister, Jonathan Dyne, Xinghao Yu, Contributing research, Liran Carmel, Khens Kermesh, Adam Bermeister, Mark Kon, Daniel Shnier

## Introduction

TP53 is a highly studied gene due to its important regulatory role in DNA, RNA (Marcel, Catez, and Diaz 2015), disease and the progression of cancer. TP53 functions as a tumor suppressor and has been described as the “Guardian of the Genome”. p53, the protein coded by TP53 predominantly binds regulatory response elements (“RE”) in introns of multiple target genes. We compared identical subsequences in introns of multiple gene/transcripts including TP53, BRCA1, men1, PELP1, SET, HIF1A, ULBP1/2 and IRF3. Some identical subsequences also contain the core RE’s of known p53 binding sites or TP53’s intronic, autoregulatory response elements (“ARE”) and strongly correlate with known miRNA’s. We describe the mechanism by which p53 concentrations change as stress free cells become adversely affected. We describe how TP53’s intronic ARE’s contribute their preference to p53 binding and propose a computational method to identify targets.

## Background

For intronic subsequences that are part of negative feedback loops (Marcel et al. 2010) steady state transcription is, in part ‘self-satisfied’ by p53 binding ARE’s. Mutant or p53 isoforms may exclusively recognize RE’s or ARE’s by iterating along DNA until identifying their uniquely compatible sequence combinations to which they bind. As soon as a stress condition affects the cell, specific p53 isoforms may be induced to relocate and bind compatible ARE’s and RE’s. More aberrant TP53 transcription may change the concentration of p53 isoforms, which would change ARE and/or RE binding patterns. Scarcity of p53 binding RE’s would also expose those genes to further unregulated or misregulated transcription events that normally include p53 and may bias transcription at other genes.

“p53 can be attenuated directly, by mutation or deletion (Herrero et al. 2016), or indirectly through alterations in methylation, miRNAs, isoform expression and p53 regulators. Six *TP53* hotspot mutations and regions potentially affected by methylation are indicated. p53 isoforms arise from the use of two alternative transcription start sites.”

“The existence in the intron of multiple identical subsequences in multiple gene/transcripts that include some or all of p53’s prerequisite [CA-><-AG] and/or [CT-><-TG] and/or [CA/CT-><-TG] (McLure 1998) core binding quarter-sites may indicate a gene/transcripts ARE or RE (Weinberg et al. 2005) state of readiness to bind p53 more completely. It should be noted the ARE or RE quarter sites and variations (Bian and Sun 1997) are each capable of binding at least one p53 monomer and stronger binding as dimers or tetramers to half and full sites has a tendency to bend DNA and increase affinity.”

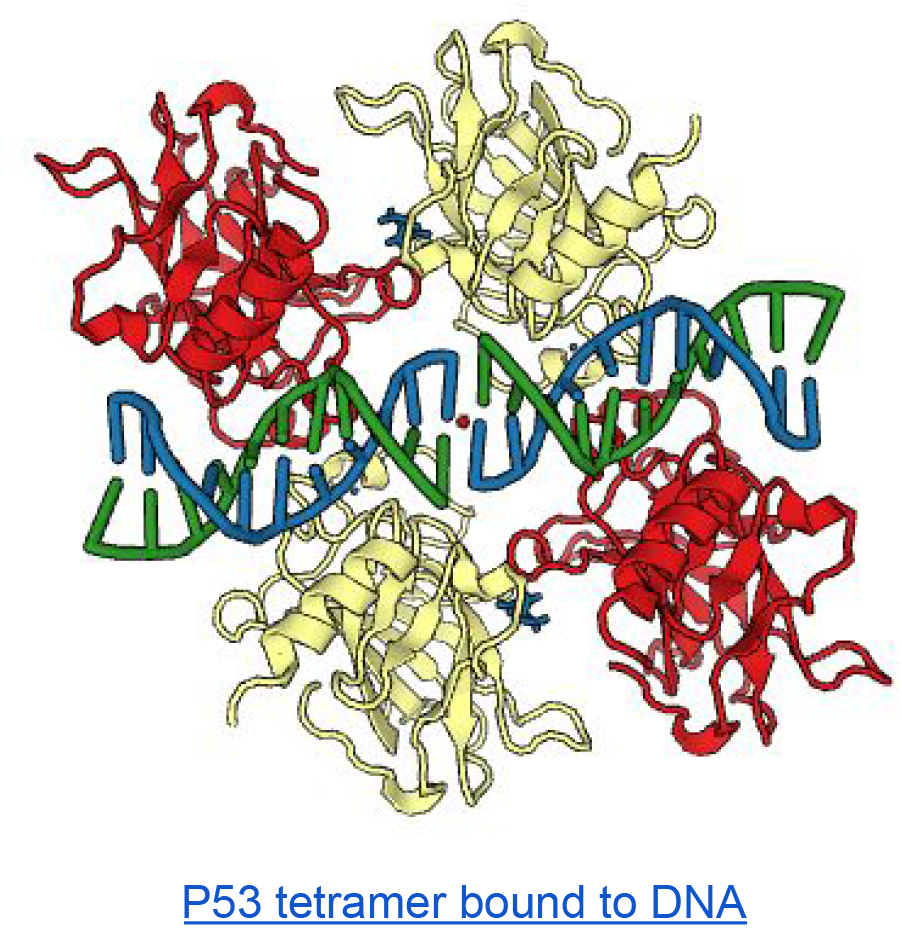

## Mechanisms of p53 Regulation in Disease

At first we loosely searched identical subsequences of multiple gene transcripts that included [CA and/or AG|CT and TG] and/or [CA and CT|TG] where each element [nucleotide pair] was separated by at least one nucleotide. We discovered the intersection of TP53 introns with other genes introns in our database. The lengths of these identical subsequences varied from 8 to more than 55 nucleotides. It is known that the regulatory regions of the most-highly regulated miRNA species are enriched in conserved binding sites for p53 (Hattori et al. 2014). Evidently the discovery, by querying the more frequently recurring quarter and half sites of ARE’s and RE’s in introns with identical subsequences confirmed that among those longer than 23 nucleotides the significant majority were known miRNAs.

Further, p53 induced (Loayza-Puch et al. 2013) transcription at combinatorial ARE-RE identical subsequences may increase cellular concentrations of these miRNA sequences more rapidly. We consider that increased concentration of ARE-RE subsequences that are capable of maturing as miRNA’s suggests they can be present in larger concentrations in regions of the cytoplasm. We found evidence in support that a fraction of the proportion of a given miRNA from an intron is capable of adjusting concentration dynamics such that it affects protein assembly and/or function:

> The first relates to a paper on Cytoplasmic Retained Introns (CIRT’s) proposing that a small fraction of the localized concentration can tip the balance. “Another important aspect of this CIRT-related phenotypic change is the number of KCNMA1 transcripts retaining i16 that can be detected in cells. Approximately 90% of detectable KCNMA transcripts in the cytoplasm are fully spliced, and 10% of the total KCNMA1 transcript population is comprised of KCNMA1i16 CIRTs. This raises the question of the physiological relevance of a relatively small number of KCNMA1 transcripts.” (Buckley et al. 2014)
>
> The second relates to the mechanism of genes in Amyotrophic lateral sclerosis (ALS) that causes concentrations to promote phase separation. “Phase transition by RNA-binding proteins (Taylor, Brown, and Cleveland 2016) that contain low-complexity domains is exquisitely dependent on concentration, and it is probable that the increased accumulation of FUS and hnRNP A1 in the cytoplasm as a consequence of mutations that affect the nuclear localization signals of these proteins is sufficient to drive excess phase separation, as shown by the hyper-assembly of stress granules in cells derived from people with relevant mutations.”

## Relative Concentration Dynamics - Codondex Computation

Concentration influences (Yu et al. 2016) DNA-Protein-RNA-Protein interactions in both the nucleus and cytoplasm, which can be significant. Further, it is evident that extremely small changes, often in a small fraction of concentrations that results from dosage sensitive genes (Babu et al. 2011) can be immediately significant.

Analogous to oil droplets in water, subcellular concentrations (Smith et al. 2016) are modified by dynamic elements including DNA, RNA, protein, metabolites, molecules and water. This concentrated environment is determinative for binding of particular elements as is the capacity for ligands to prevail in the pressure of condensed space. The effects on intrinsic protein disorder (Babu et al. 2011), including p53 is also concentration dependent and dosage sensitive. Double stranded DNA association and dissociation inherencies, as influenced by epigenetic factors and induced by proteins (Pontius and Berg 1992) may play a role in organizing (Elbarbary, Lucas, and Maquat 2016) DNA subsequence concentrations to be more or less favorable to binding protein, transcription efficiency and translation.

Intra chromosome, subsequences of the DNA molecule can be thought of as concentrations. Each gene is then a particular pattern of subsequence concentrations that relatively and proximally define DNA of the chromosome. Therefore, our interest in specifics of DNA’s relative concentrations led us to review every adjacent combination of any of the four letters A,G,C or T within a gene of interest. Our DNA kmer algorithm computes and maintains a database of every possible subsequence >7 letters of a given DNA sequence. It captures the smallest single nucleotide change in any subsequence >7 for all possible subsequences of a transcript.

By example, from the transcribed DNA sequence GAGCTTCGAG_6_ there is potential to momentarily exist, including as derivative RNA subsequences or sub-concentrations >7 |GAGCTTCG_3_| |AGCTTCGA_4_| GAGCTTCGA_4_| GCTTCGAG_3_| AGCTTCGAG_4_|. Subscript represents the repeat frequency each subsequence in any >7 derivations of GAGCTTCGAG. In other words if two subsequences have the same repeat frequency they will have the same relative concentration and will likely be subjected to similar dissociation dynamics resulting from transcription events. Since DNA nucleotide order is tightly maintained this model also represents the complete weighted set of RNA subsequences that can be potentially transcribed into the nucleus and cytoplasm. Because various concentrations of repeatedly transcribed DNA will exist as RNA in the cell at various times, we devised an exhaustive model in which a single cell-genes’ ‘any-time’ potential state is computed. Measuring and sequencing a genes’ cytoplasmic RNA concentrations and comparing them to the any-time potential state in same cell transcripts may provide a useful comparative. Using this as a diagnostic tool may also provide a sensitive holistic measure to associate non-coding DNA-RNA states with diseases.

As discussed in this paper, RNA in various forms may bind protein and in some cases cause protein to relocate to regions where it interacts. These types of reactions are abundant in concentrations, which are sometimes determinative to binding and thereafter to downstream reactions. We looked at the intersection of TP53 and IRF3 supported by their known interactions; The novel findings presented by Baresova and colleagues (Baresova et al. 2013) are that “vIRF-3 can significantly decrease the stability of p53 protein, its phosphorylation, tetramer formation, and DNA-binding ability. This results in reduced transcription of the p53-regulated genes, the p21 and Bax genes, which may lead to impaired cell growth or apoptosis. These data indicate that Kaposi’s sarcoma-associated herpesvirus evolved mechanisms to attenuate the p53 growth-regulatory pathway and thus contributes to development of virus-associated malignancies.”

## Method

For any intron, we partition DNA transcript data by its computed, potential subsequences (potential kmers “pkmers”) associated with a signature of mRNA or protein encoded by the transcript. For each of multiple same gene transcripts, pkmers are computed by iterating single nucleotide steps to establish every potential subsequence in our database. Once complete for the entire intron, we count repeats (within the intron computation) for each pkmer and compute rankings. We assimilate ranked results from each step of each transcript-intron to a highly structured multi-transcript, multi-dimensional vector.

For any given pkmer (which we may refer to by offset#) we generate an intra transcript-intron sequence ranking. We compute each pkmers’ iScore, derived by summing trailing (3’) repeats found in it, which we divide by the pkmer length. The protein signature of every pkmer is constant for the transcript-intron. Any step (next offset#) of the transcript can be compared using the vector comprising iScore:Protein signature as assimilated for all transcripts.

The illustration below is for offset#48333 of 15 transcripts of men1. Each Transcript ID, iScore and Protein Rank Position is recorded in a vector (the illustration subscripts last 3 digits of transcript ID). Since protein signature is constant for every pkmer of a given transcript-intron for a gene, order by the protein signature is static (“PhV”). Rules permit shuffling transcripts offset#’s that share the same iScore, as per transcripts 422 and 326 to maximize transcript assimilation in the vector. Finally we assign vector positions 1-15 to transcripts according to their order, this constitutes Position in Vector (“PiV”).

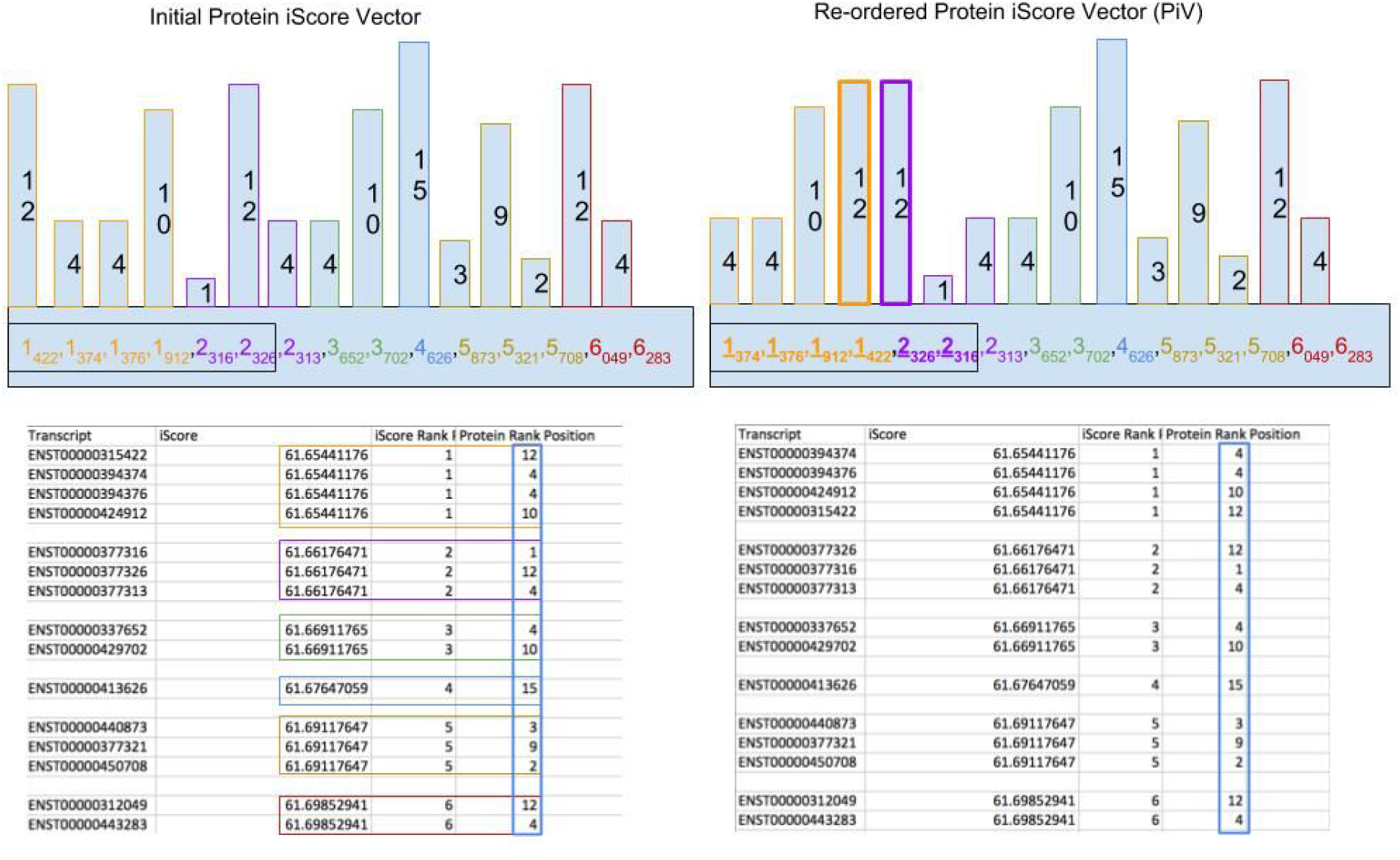

By example, a computation of 50 nucleotides of each transcript-intron subsequence amplifies to ±4000 rows (steps). Each row contains one of all possible pkmers for the total length 50 intron sequence. We sort rows by pkmer aggregate repeats and identify pkmers that contain the most changes in the vector order at the next row. The vector order changes because of a transcripts iScore:Protein signature assimilation to the vector for each offset#.

Membership to the vector is bounded by nucleotide order of each transcript-intron DNA and transcripts’ protein signature. Therefore, a change in a transcript PiV from step (row) to step (row) must be understood differently to the row with the most changes in all transcript PiV orderings. The latter being the biggest PiV change, by repeats for all transcript-intron subsequences for the set of pkmer:Protein assimilations.

The result is a selection method, for any given sequence length, from all pkmers of a set of transcripts. The identification is the pkmers that most strongly modified PiV in a successive step when sorted by aggregate pkmer repeats.

## TP53 intronic half-sites implicate p53

We demonstrate that TP53 quarter and half sites implicate p53 according to Codondex iScore-PiV computations. Introns of TP53 reveal known ARE half-sites 5’-PuPuPuC(A/T)|(A/T)GPyPyPy-3’ for the p53 dimer or tetramer that infer its four nucleotide core as a strong signal for binding preference. The core combinations CAAG, CATG, CTAG or CTTG in the 10 letter ARE or RE context are sufficient to ensure high degrees of half-site binding specificity. Further, p53 dimers can (McLure 1998) bind reversed, adjacent quarter sites as a reverse half site core 5’-G(A/T)|(A/T)C-3’ - GAAC, GATC, GTAC or GTTC, which interestingly we discovered are strongly implicated in men1.

Analogous to setting a combination in a lock then unlocking-locking using the combination: We investigated whether TP53 intron-1 ARE’s alter p53 protein through transcription-translation (Gabunilas and Chanfreau 2016) or post-translational modification (Gu and Roeder 1997) in a way that would transmit to binding domains of p53 monomers (Tafvizi et al. 2011) a specific ARE (or RE) preference. One way may be through DNA to by the mechanism of p53 successfully binding intronic and exonic (Yang and Wu 2004) RE’s of genes to form tetramers and induce transcription. Further we considered how this may implicate RE’s in other genes p53 is known to bind. For RE’s we reviewed identical pkmers (>7 letters) of TP53 transcripts with other gene pkmers that also contain the core binding domain. We queried all identical pkmers with any six (6) letter (1-4-1) half-site cores. We initially used “GCAAGC” and discovered several genes that shared identical pkmers with this 6 letter core including HIF1A with TP53. We queried the identical subsequence cores on multiple gene intron subsequences and discovered their intersections with half-site cores in RE’s or ARE’s.

In summary, monomer-dimer-tetramer binding RE-ARE’s suggests their innate sensitivities are reflected in their determinative structures. Binding events are ultimately governed by the relationship between ligand and DNA sequence, which are influenced by complex factors that diminish the chance a tetramer will form and through which transcription will be promoted. This presents a very wide range of statistical outcomes with dependencies in ligand, RE-ARE sequences, upstream and downstream DNA or RNA concentrations and various epigenetic factors.

The image below from Excel file was ordered by repeats (column O) and pkmer length. We searched all pkmers containing AGGCAAGCCC - one of four specific half-site binding sequences of interest and used example TP53 ENST00000420246 @offset:#512580 [Start 1010,End 1019]. We highlighted (light green) every pkmer (column G) that contained the full pkmer of offset #512580 and in continuous light green all offsets# that start at nucleotide #1010 of the intron - column H. We observed changes in the vector order (column E) as implicated by repeats and reverse complement (“coffset”-column R).

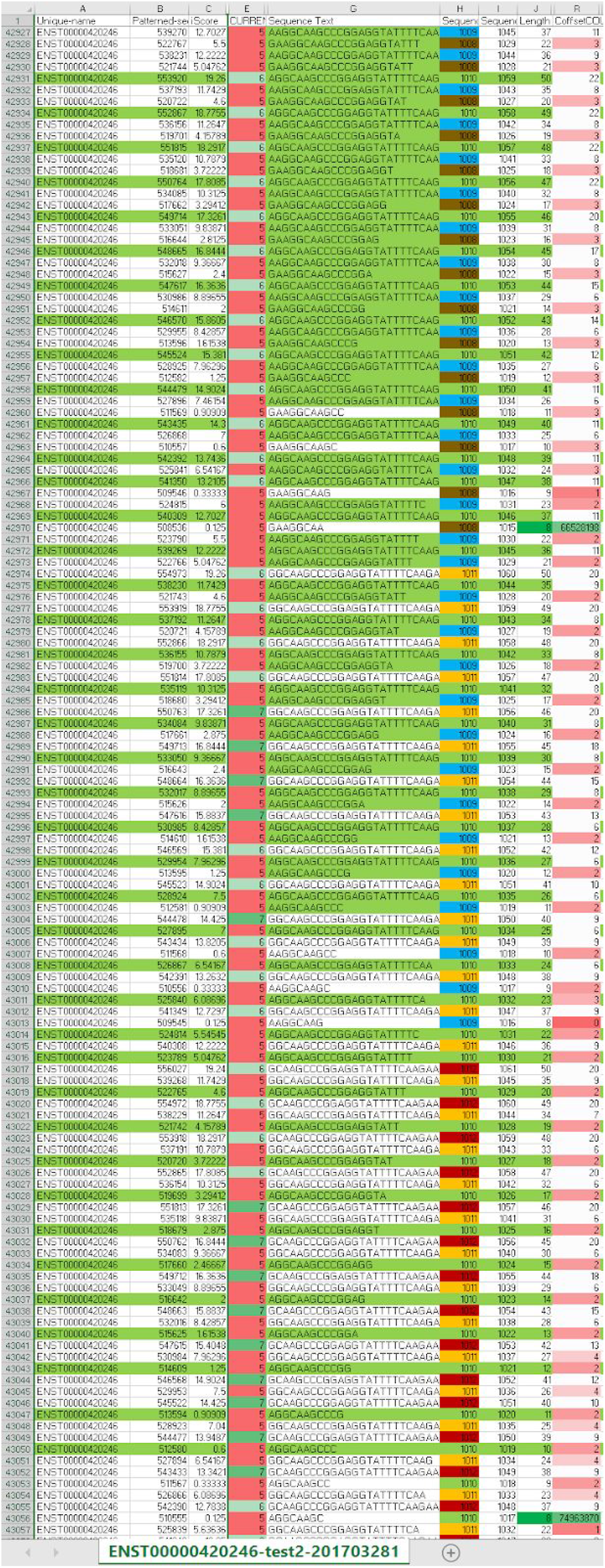

The suggested relative concentration score of each pkmer is represented by repeats (column O). The changes in pkmers PiV at each progressive step suggest the TP53 transcript-intron transmits modification to its associated p53 monomer (Dornan et al. 2003). This may be sufficient to identify subsequences that infer modification to p53 (and ligand) to signal specific transcription outcomes to specific binding combinations of ARE’s or RE’s that promote transcription of RNA derivatives.

## ID Gene-Transcript-Intron-Protein Subsequences

We searched all transcript-intron computations to discover the shortest length pkmer that also included the highest number of length 8 pkmers associated with the biggest change in PiV at the next step (row). We considered for any given pkmer length, that the highest number of PiV changes (biggest total vector shifts) would be of interest. The leading candidate was ENST00000359597.

To verify PiV variations we reviewed (image below) ENST00000359597 (left) against two randomized versions using a transcript wide random letter reordering (middle) and separately a random pick A.C.T.G of each nucleotide (right). The orange bulge confirmed best and most consistent, per offset# correlation of PiV, coffset and PiV shifts occurred in the ENST00000359597 iScore report (left).

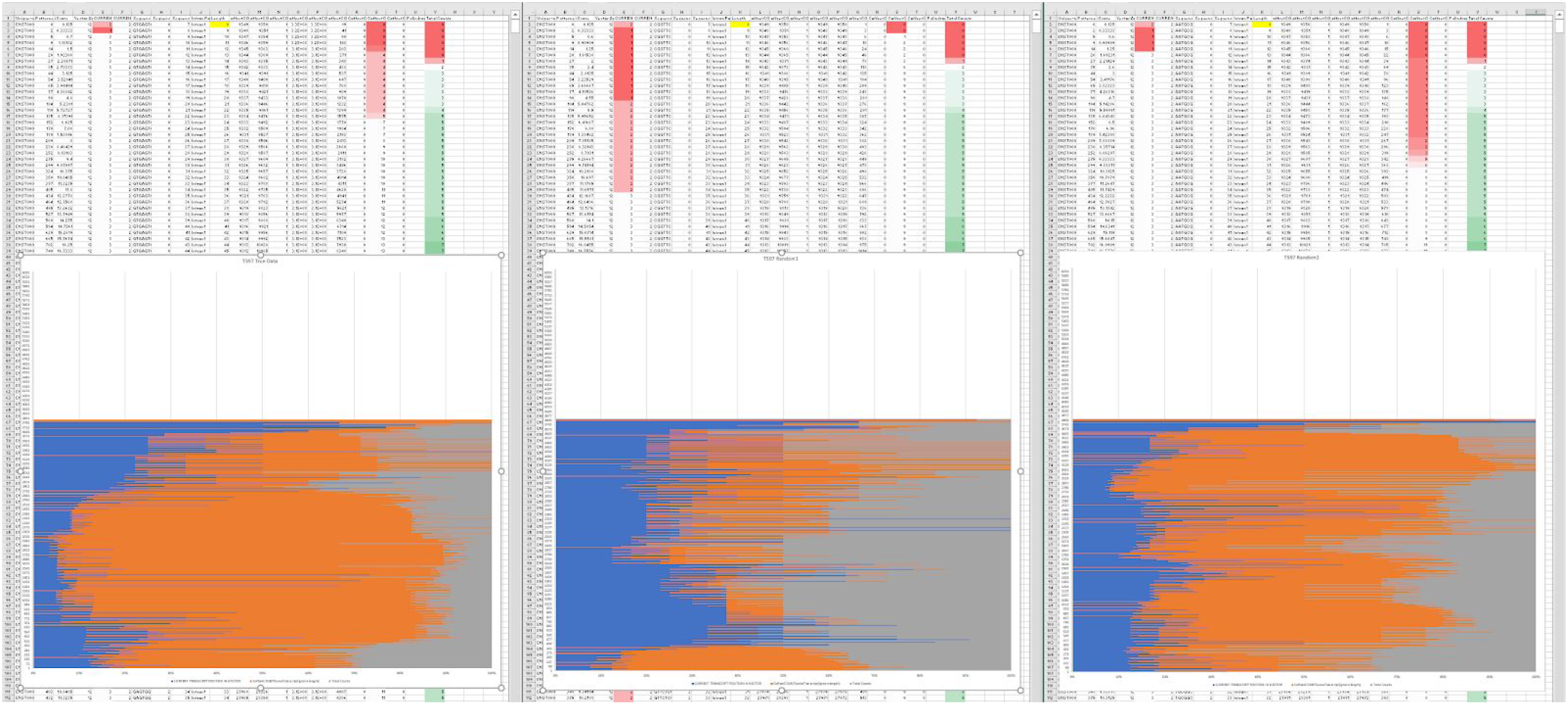

We then identified the frequency that length 8 pkmers with biggest PiV shifts were incorporated in their length 50 pkmers that shared the same sequence start nucleotide in the intron;

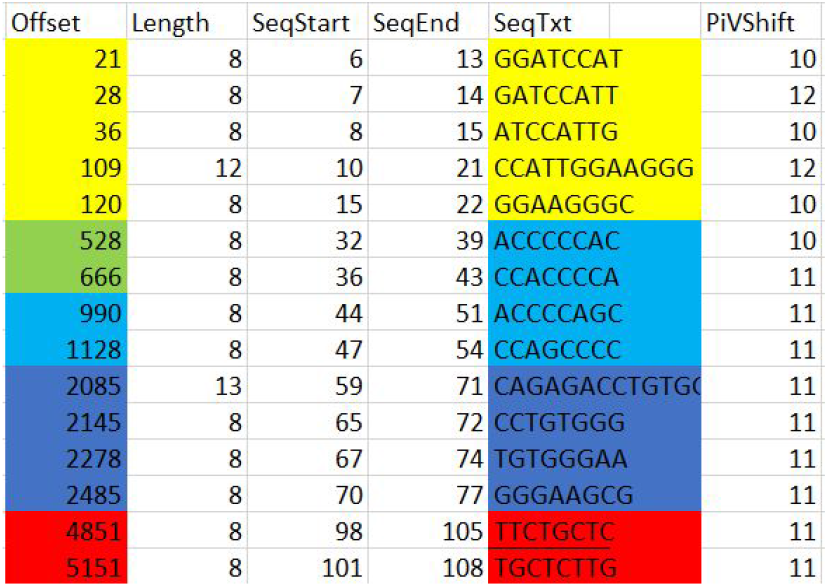

As seen above, these sequences are grouped according to their intron DNA sequence start (SeqStart) positions and are overlapping. We were able to immediately identify using offset#21 that its length 50 counterpart at offset#1218 (below) contained 9 other offsets from the above group (_subscript_ below identifies SeqEnd for each pkmer in the table above).

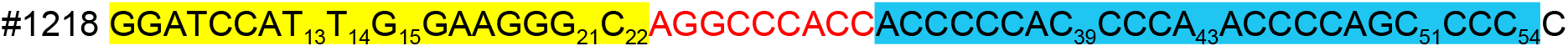

We then confirmed (below) that the identical sequence data, referred to earlier in this paper contains ENST00000359597, offset#21 and although not a classical p53 binding site, it conforms with lesser binding site rules.

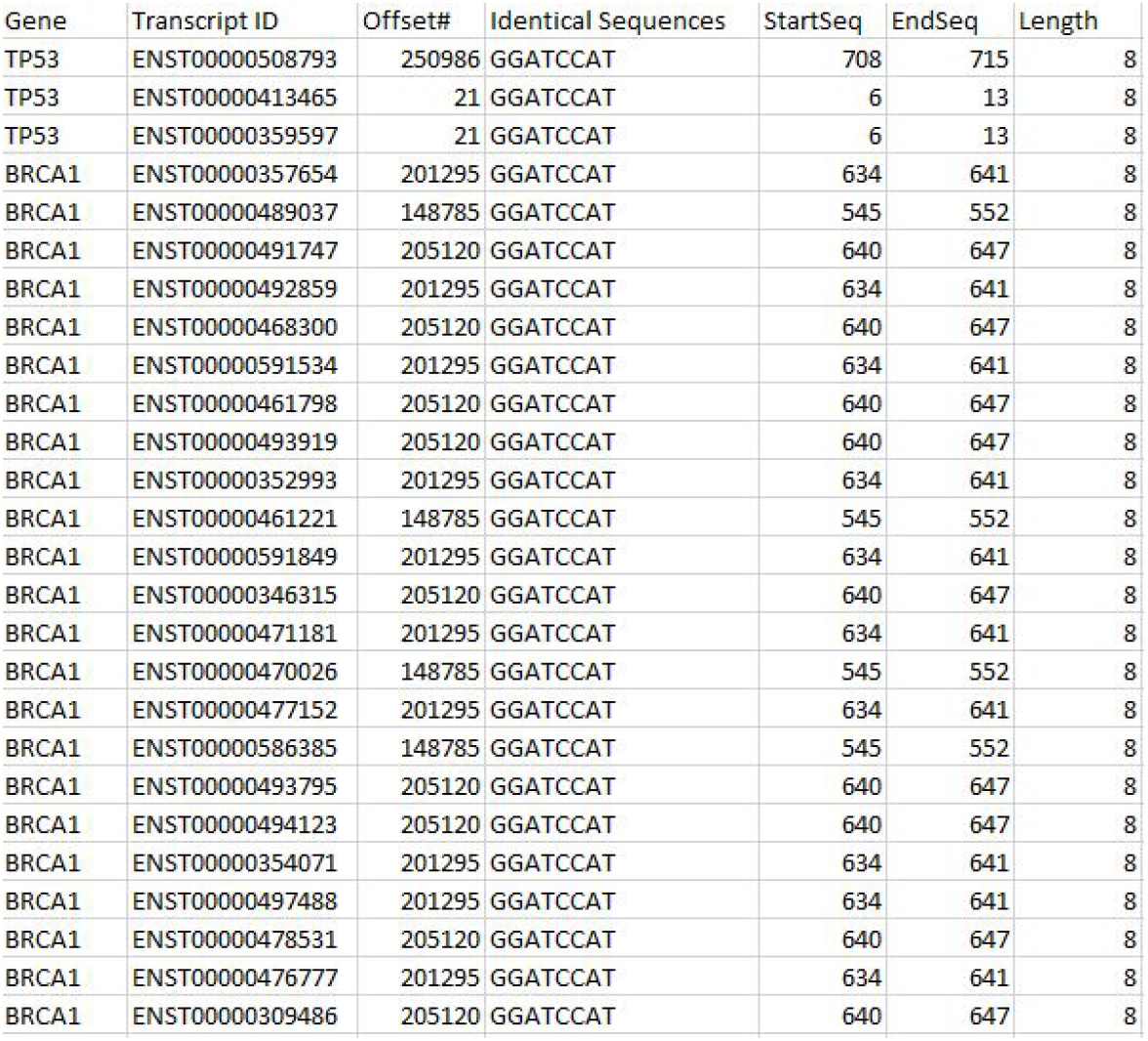

We further confirmed that the ENST00000420246 sequence at offset #21 (below) is also an identical sequence together with several other TP53 transcripts.

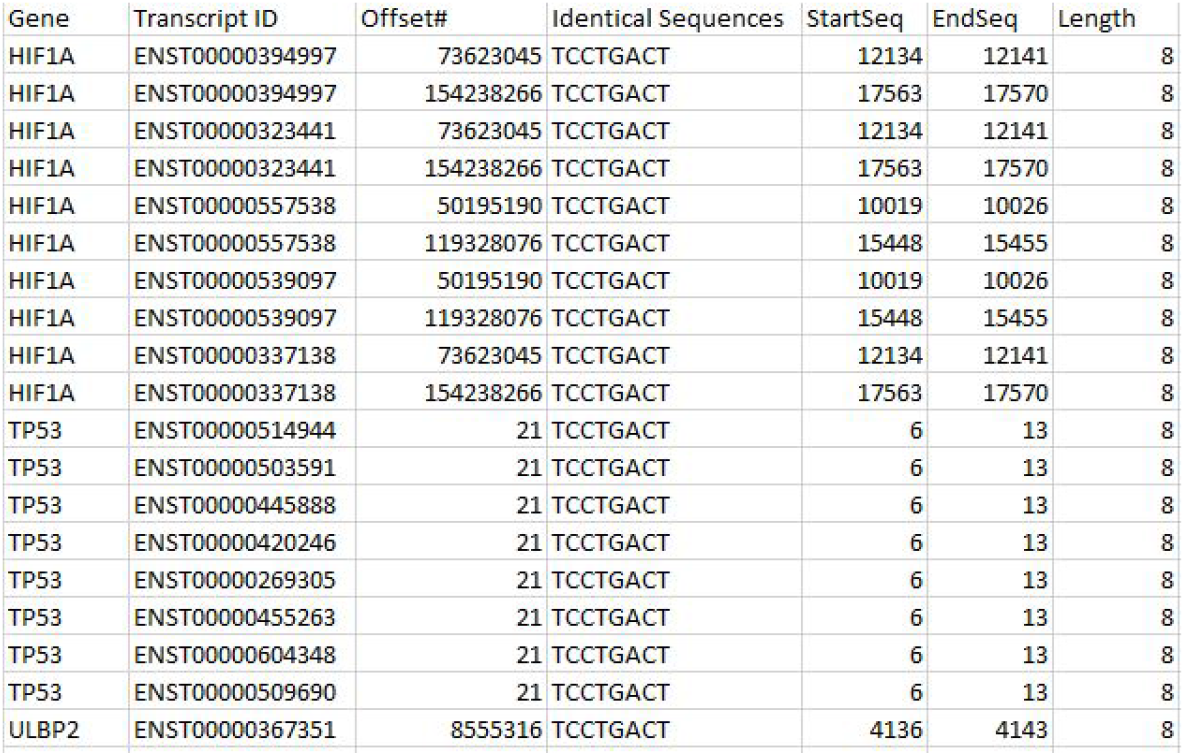

We confirmed in P53 that the length 8 offset# associated with the set of highest scoring transcript-wide PiV position shifts will have a SeqStart where its Length(8+n) will also contain the most other length 8 offset subsequences of the set.

The discovery of gene-wide, multi-transcript, shortest length offsets containing the most number of biggest PiV shifts is a novel way to discover relevant intronic subsequences. Further testing of these subsequences in biology will confirm whether their inherent, relative concentration makes them capable of affecting p53 or more specifically binding performance of the p53 monomer as suggested in this paper.

## Conclusion

Codondex is a novel multi-transcript, single cell analysis of inherent and relative concentration classifications in DNA. It is assembled by DNA/intron pkmers to identify heritable effects on protein by comparing modifications to the order of transcripts in the PiV, protein vector. It is proposed that concentration change induces different combinations of p53, according to their inherited traits to translocate where they bind highly specific ARE’s or RE’s, which are subsequently transcribed. Further work is required to identify TP53 ARE’s and determine their capacity to modify p53 ligand or binding conformation through specific concentration changes the transcribed TP53 feature induced.

